# BepiPred-3.0: Improved B-cell epitope prediction using protein language models

**DOI:** 10.1101/2022.07.11.499418

**Authors:** Joakim Clifford, Magnus Haraldson Høie, Morten Nielsen, Sebastian Deleuran, Bjoern Peters, Paolo Marcatili

## Abstract

B-cell epitope prediction tools are of great medical and commercial interest due to their practical applications in vaccine development. The introduction of protein language models (LM) trained on unprecedented large datasets of protein sequences and structures, tap into a powerful numeric representation that can be exploited to accurately predict local and global protein structural features from amino acid sequences only. In this paper, we present BepiPred 3.0, a sequence-based epitope prediction tool that, by exploiting LM embeddings, greatly improves the prediction accuracy for both linear and conformational epitope prediction on several independent test sets. Furthermore, by carefully selecting additional input variables and epitope residue annotation strategy, performance can be further improved, thus achieving extraordinary results. Our tool can predict epitopes across hundreds of sequences in mere minutes. It is freely available as a web server with a user-friendly interface to navigate the results, as well as a standalone downloadable package.

## 1 Introduction

B-cells are a major component of the adaptive immune system, as they support long-term immunological protection against pathogens and cancerous cells. Their activation relies on the interaction between specialized receptors known as B-cell receptors (BCRs) and their pathogenic targets, also known as antigens. Upon interaction, B-cells produce antigen-specific molecules known as antibodies, which are identical to BCRs in structure, except that they do not have a transmembrane region. More specifically, BCRs selectively interact with specific portions of their antigens known as epitopes. B-cell epitopes are divided into two types. Linear epitopes are found sequentially along the amino-acid sequence, while conformational epitopes are interspersed in the antigen’s primary structure, and brought together in spatial proximity by the antigen’s folding. While approximately 90% of B-cell epitopes fall into the conformational category, most of these contain at least a few sequential residue stretches [14]. Epitopes are typically found in solvent-exposed regions of antigens. Physical and chemical features other than solvent accessibility, such as hydrophobicity, secondary structure propension, protrusion indexes, and local amino acid composition, have been shown to affect the likelihood of epitopes [14]. B-cell epitope identification is of great interest in biotechnological and clinical applications, such as attenuated or subunit vaccine designs and therapeutic antibody development. Their identification, however, is a costly and time-consuming process requiring extensive experimental assay screening. In silico prediction methods can significantly reduce identification workloads by predicting epitope regions, and because of this they have become critical for such tasks [20][19]. Structure-based tools have been developed for predicting B-cell epitopes [15][1][25][24]. However, as experimentally determined structural information is often not available, epitope identification must in many cases be performed from amino-acid sequences alone. So far, sequence-based tools have only achieved mediocre results, in general worse than structure-based tools [21][18][28][9]. Thanks to recent developments in the field of machine learning, models trained on large datasets of protein sequences and structures are now available to accurately predict local and global protein structural features from amino acid sequences only [10][8]. In particular, protein language models (LM) have been demonstrated to allow for a powerful numeric representation of protein sequences, that in turn can be exploited to substantially increase the accuracy in many different prediction tasks [17][4]. Here, we present BepiPred-3.0, a sequence-based tool, which utilizes numerical representations from the protein language model ESM-1b, to vastly improve prediction accuracy for linear and conformational B-cell epitope prediction [[17]. Furthermore, by carefully selecting the architecture of the predictor, the training strategy, additional input variables to the model, and using an epitope residue annotation strategy adopted from one of our earlier works, performance can be further improved, thus achieving unprecedented results [26].

## 2 Methods

### 2.1 Structural datasets

A first dataset, named BP2, consists of the antigens used for training the BepiPred 2.0 server. This dataset contains 776 antigens and is available at the BepiPred-2.0 web server. A second updated dataset, named BP3, was built using the same approach previously adopted in BepiPred 2.0 [9]. We first identified crystal structures from the Protein Data Bank deposited before 29/09/2021 that contain at least a complete antibody, and at least a non-antibody (antigen) protein chain [3]. This was done using existing HMM profiles developed in-house [12]. We only included crystal structures with a resolution lower than 3 Å and R-factor lower than 0.3. On both datasets, we identified epitope residues using the same approach adopted in our previous paper [9]. On each antigen chain, we labeled every residue that had at least one heavy atom (main-chain or side-chain) at a distance of less than 4 Å to any heavy atom belonging to residues in antibody chains of the same crystal structure as an epitope residue. We only retained antigen chains with at least one epitope residue and with a minimum sequence length of 39. Epitope residues for antigens which were 100% identical in sequence were merged and only one antigen entry included. Missing residues were not included as part of the epitope annotated antigen chains. After these steps, we obtained a total of 1466 antigens for the updated BP3 dataset.

### 2.2 Redundancy reduction

We used a redundancy reduction approach similar to existing works, which we called the epitope collapse strategy [26]. Here, sequence clusters were first generated using MMseqs2 [22]. Next, all antigen sequences belonging to the same cluster were aligned to the MMseqs2 defined cluster representative. The cluster representative sequence then underwent the following modifications: At any position of the alignment, if an epitope was identified in the cluster representative, it was retained as is. If an epitope was found on any of the aligned antigen sequences, it was grafted onto the cluster representative sequence and labelled as epitope. For each cluster, only the sequence of the cluster representative, modified as described above, was retained. This was done at 95% sequence identity for both the BP2 and BP3 datasets, making the size of the BP2 and BP3 datasets 238 and 603 antigens, reduced from 776 and 1466 antigens respectively. Furthermore, the strategy was performed at 50% sequence identity for the BP3 dataset. This dataset was called BP3C50ID and contained 358 antigens reduced from 1466 antigens.

Redundancy was further reduced for two additional datasets. The sequences of the final BP2 and BP3 datasets described above were clustered at 70% sequence identity using MMseqs2, and then only the cluster representatives were incorporated into the reduced datasets [22]. This gave rise to datasets BP2HR and BP3HR, which contained 190 and 398 antigens, reduced from 238 and 603 antigens, respectively (Table 1).

**Table 1:**
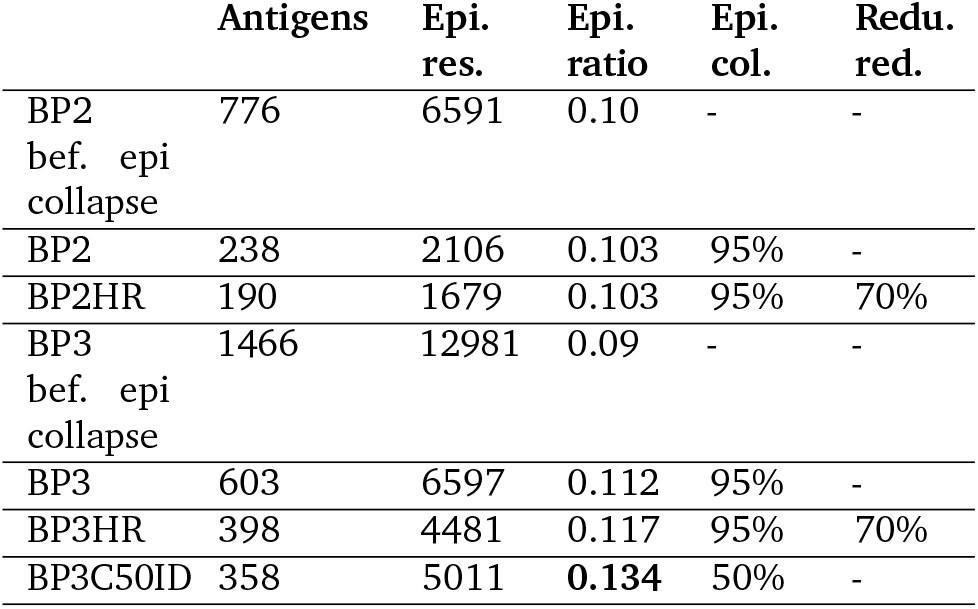
The number of antigens, epitope residues (**Epi. res**.), epitope residue ratios (**Epi. ratio**) as well as redundancy reduction (**Redu. red**.) and epitope collapse (**Epi. col**.) by MMseqs2 for BP2, BP3, BP2HR, BP3HR and BP3C50ID. Epitope residue ratios were computed as the ratios between the number of epitope residues and total number of residues. The redundancy reduction and epitope collapse columns are the MMseqs2 sequence identity %’s, for which the epitope collapse and the second redundancy reduction approaches were used.

### 2.3 External test sets

Three different independent test sets were built. The first corresponds to the same 5 antigens used as external test evaluation in BepiPred-2.0 [9]. But the antigens were extracted from BP3C50ID, and therefore contained updated and enriched epitope annotations. Any sequence with more than 20% sequence identity to this test set, as calculated by MMseqs2, was removed from all BP2 training sets. After this removal, BP2 and BP2HR training sets contained 233 and 185 antigens. The second external test set comprises the antigens mentioned above, plus 10 additional BP3C50ID antigens selected from the MMseqs2 clusters at 20% identity. Here, any sequence with more than 20% identity to any of the 15 antigens was removed from all BP3 training sets. After this removal, the training sets for BP3, BP3HR and BP3C50ID contained 582, 383 and 343 antigens. Finally, a third external dataset was constructed by downloading all linear B cell epitopes from the IEDB and then discarding any epitopes containing post translational modifications, as well as epitopes for which the source protein ID, as indicated in the IEDB entry itself, was not a UniProt entry [27][2]. Epitopes with a perfect match in the source protein were mapped to the relevant region while the rest were discarded. This resulted in 4072 epitopes with mapped protein sequences. Finally, we removed all proteins which had more than 20% sequence identity to the BP3C50ID dataset, leaving 3560 sequences (Table 2).

**Table 2:** The number of antigens, epitope residues (**Epi. res**.), epitope residue ratios (**Epi. ratio**) and epitope collapse strategy (**Epi. col**.) for the 5 and 15 antigen external sets as well as the external IEDB test set. Epitope residue ratios were computed as the ratios between the number of epitope residues and total number of residues. The epitope collapse strategy column is the MMseqs2 sequence identity %, for which epitope collapse was done.

|  | Antigens | Epi. res. | Epi. ratio | Epi. col. |
| --- | --- | --- | --- | --- |
| External test set 1 | 5 | 88 | 0.256 | 50% |
| External test set 2 | 15 | 209 | 0.190 | 50% |
| External IEDB test set | 3560 | 8818 | 0.116 | - |

### 2.4 Dataset encoding

Residues were represented either by sparse encoding, BLOSUM62 log-odds scores or by numeric embeddings extracted from the ESM-1b protein language model [17]. When employing sparse and BLOSUM62 encodings, the encoding of each residue was generated by concatenating sparse or BLOSUM62 encodings from the residue itself and from the 8 neighboring residues, 4 on each side. For residues close to sequence terminals where an insufficient number of neighbors existed, padding tokens were used to fill in for lacking residues. Thus, each residue was represented by a vector of size 189 (9*21). ESM-1b encodings were obtained by passing antigen sequences through the pretrained ESM-1b transformer, and extracting the resulting sequence representations from the model. Here, each residue was represented by a vector of size 1280. We also included the NetSurfP-3.0 predicted RSA values for the central residue and the protein sequence lengths to the encodings [8]. The target values were encoded in a position-wise binary manner, resulting in a sequence denoting epitope and non-epitope residues with the same length of the antigen sequence (Figure 1).

**Figure 1:**
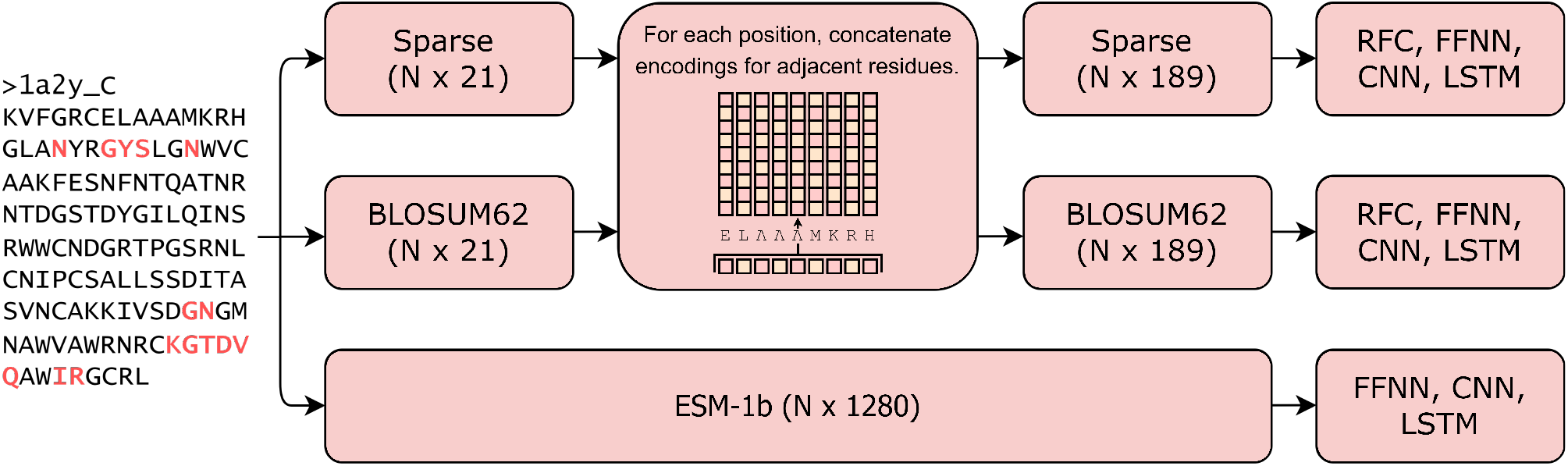
Overview of sequence encoding pipelines, where N is length of the sequence. Amino acid sequences were encoded using either sparse, BLOSUM62 or ESM-1b derived sequence representation schemes. For the two former approaches, encodings from adjacent residues were concatenated to generate a new set of encodings describing the sequence context of each residue. The encoded sequences were subsequently used for training various models for position-wise antigen prediction.

### 2.5 Model architectures and hyperparameter tuning

Feed Forward (FFNN), Convolutional (CNN), and Long Short-term Memory (LSTM) neural networks were trained on sparse, BLOSUM62 and ESM-1b encodings, with or without the additional variables (sequence length or NetSurfP-3.0 predicted RSA values). Model weights were initialized and updated using the default PyTorch weight initialization schemes and an Adam optimizer [11]. Since we have independent test data, hyperparameter tuning could be performed in a simple 5-fold cross fold validation setup using a grid search on the training sets described in method section 2.3. Hyper-parameters were chosen as those which yielded the best validation cross-entropy (CE) loss averaged across all folds. The hyperparameters in question are the learning rate, weight decay, dropout rate and different architectural setups (see supplementary methods section 5.3 for the exact final model configurations). As a baseline, a Random Forest Classifier (RFC) was trained on sparse and BLOSUM62 encodings. We tested forest sizes ranging from 25-300 and determined the optimal size on a validation AUC score basis (see supplementary method section 5.2).

### 2.6 Training and evaluation

A 5-fold cross-validation was used to train the models with the optimized hyperparameters discussed in the previous section. For BP2 and BP3 cross-validation setups, antigens were clustered at 70% sequence identity using MMseqs2, and sequences of the same cluster placed into the same partition. A total a five partitions were created [22]. For the 3 remaining datasets, training and validation splits were made randomly. Each cross-validation setup generated 5 models, where both validation cross entropy (CE) loss and AUC was used as an early stopping criteria. For instance, the training procedure for a fold was the following: Model weights were initialized using the default PyTorch weight initialization schemes. The model was trained on the training data using batches of 4 antigens, and the loss backpropagated using the Adam optimizer [11]. After each epoch, a cross entropy loss and AUC score was computed on the validation set. Only if both scores improved, were the model parameters stored. We trained for 75 epochs. Next, the models were evaluated on the independent test data sets. The evaluative metrics on the external test set were AUC, AUC10, MCC, recall, precision, F1-score and accuracy. AUC scores were computed by concatenating all 5 model outputs from all 5 model outputs, into a single vector, and comparing with a 5 times duplicated label vector. AUC10 scores were computed as the integral of the ROC curve area going from 0 to 0.1 on the false positive rate axis, divided by 0.1, setting the AUC10 score in a range of 0 to 1. For the remaining threshold-dependent metrics, a majority voting scheme was used based on the individual predictions from the 5 models. The classification threshold used for each fold model was one that maximized the MCC score on respective validation splits. For some model performance comparisons, paired t-tests were performed and p-values calculated on their CE loss scores on all test antigen residues.

## 3 Results

### 3.1 Improved performance on BP3 datasets and when the epitope collapse strategy is used

In an initial benchmark study, we investigated the effects of using updated datasets (BP3, BP3HR and BP3C50ID) versus datasets constructed from the BepiPred2 paper (BP2, BP3HR), as well as various redundancy reduction approaches (see method section 2.2 for more details). We determined that BP3 datasets (BP3 and BP3HR) led to a improved predictive performance compared to models trained using the smaller BP2 datasets (BP2 and BP2HR). This was observed by training a set of random forest classifiers (RFCs) and evaluating on the 5 antigen external test set. We improved performance further by first doing sequence redundancy reduction at a 50% identity threshold as defined by MMseqs2, and then using the epitope collapse strategy (BP3C50ID dataset) where all epitopes for sequences found in a given cluster are transferred to the cluster representative antigen. We determined that the performance increases from using updated datasets and the epitope collapse strategy, were statistically significant at all common thresholds (p-values of 1 × 10^*−*15^ and 1 × 10^*−*20^ respectively) (for more details on the epitope collapse strategy refer methods section 2.2, and for details on the results, see supplementary result section 6.1). Given these results, we therefore only used the BP3 and BP3C50ID datasets for the subsequent analyses.

### 3.2 Improved performance using neural networks and ESM-1b sequence embeddings

The main goal of this paper was to demonstrate that a B-cell epitope predictive tool based on LM embeddings will perform better than models using other encoding schemes, such as BLOSUM62 or sparse encoding. To assess the best performing machine-learning architecture and sequence representation for the BP3 and BPC50ID datasets, optimal hyperparameters for four different architectures (RFCs, FFNNs, CNNs and LSTMs) were identified using a grid search (see methods section 2.5). For each architecture, we investigated the performance when representing protein residues with sparse encoding, BLOSUM62 encoding or ESM-1b embedding. The best performing models were selected from optimal cross-validation performance, and tested on the 15 antigen external test set using a classification threshold optimized on the validation splits (Table 3). We found that all neural networks using ESM-1b sequence embeddings as input performed better. We determined that the performance increase of models using ESM-1b embed-dings instead of BLOSUM62 encodings was statistically significant at all common thresholds (p-values of 1 × 10^*−*39^, 1 × 10^*−*16^ and 1 × 10^*−*38^ for the FFNN, CNN and LSTM respectively). The paired t-test was performed by comparing each models CE loss scores on all test antigen residues. The overall best performing model was the FFNN architecture using ESM-1b sequence embeddings as input. Contrary to our expectations, the FFNN (ESM-1b) performed better than the CNN (ESM-1b) and LSTM (ESM-1b) models. While CNNs and LSTMs sequentially process residues along the antigen sequence, the FFNN was trained on single residue ESM-1b encodings without applying any sliding window to the input data. This suggests that the ESM-1b residue embeddings already contain sufficient information about the sequence neighborhood, making convolutions over the sequence unnecessary. We also did an identical analysis for models trained on the BP3 dataset, which gave us slightly worse results than those presented in Table 3 (see supplementary results section 6.2). And so similar to the initial benchmark using RFCs, we find that the models trained on the BP3C50ID dataset perform better. This points to the epitope collapse strategy improving performance.

**Table 3:** Evaluation performance on an external test set of 15 antigens, with a set of RFC, FFNN, CNN and LSTM models, trained using cross-validation on the BP3C50ID dataset. The best models in terms of validation CE loss and AUC score were chosen for evaluation on the test set (see methods section 2.6). The evaluative metrics used were AUC, AUC10, MCC, recall, precision, F1-score and accuracy. The best score within a metric category is marked in bold. We also computed paired t-test p-values, comparing the CE loss scores of models using BLOSUM62 encoding and ESM-1b embeddings as input.

| BP3C50ID Models | AUC | AUC10 | MCC | Recall | Precision | F1 | Accuracy | P-value |
| --- | --- | --- | --- | --- | --- | --- | --- | --- |
| RFC (Sparse) | 0.593 | 0.091 | 0.104 | 0.535 | 0.237 | 0.328 | 0.585 | - |
| RFC (BLOSUM62) | 0.611 | 0.096 | 0.129 | 0.593 | 0.245 | 0.347 | 0.576 | - |
| FFNN (Sparse) | 0.631 | 0.104 | 0.153 | 0.716 | 0.243 | 0.363 | 0.522 | - |
| FFNN (BLOSUM62) | 0.630 | 0.103 | 0.155 | 0.730 | 0.243 | 0.364 | 0.516 | - |
| FFNN (ESM-1b) | **0.697** | **0.118** | **0.220** | 0.658 | 0.289 | **0.401** | 0.627 | $< 1 \times 10^{-39}$ |
| CNN (Sparse) | 0.635 | 0.101 | 0.154 | **0.746** | 0.240 | 0.364 | 0.504 | - |
| CNN (BLOSUM62) | 0.636 | 0.101 | 0.158 | 0.723 | 0.245 | 0.366 | 0.523 | - |
| CNN (ESM-1b) | 0.685 | 0.115 | 0.205 | 0.585 | 0.293 | 0.390 | 0.653 | $< 1 \times 10^{-16}$ |
| LSTM (Sparse) | 0.606 | 0.092 | 0.113 | 0.732 | 0.225 | 0.344 | 0.469 | - |
| LSTM (BLOSUM62) | 0.641 | 0.103 | 0.147 | 0.542 | 0.261 | 0.353 | 0.622 | - |
| LSTM (ESM-1b) | 0.685 | 0.116 | 0.214 | 0.596 | **0.297** | 0.396 | **0.655** | $< 1 \times 10^{-38}$ |
| BP3C50ID overall average | 0.641 | 0.104 | 0.159 | 0.651 | 0.256 | 0.365 | 0.568 | - |

Importantly, we also investigated how the epitope collapse strategy affects predictions on a test set without collapsed epitopes. We note that the collapsing of epitopes also includes the possibility of adding additional epitope residues to the chosen sequence, which in turn would modify the extracted ESM-1b sequence embeddings. Here, we found that there was no apparent decrease in performance (see supplementary results 6.5).

To conclude, we find that while part of the improvement can be ascribed to training on a larger dataset and the epitope collapse strategy (see result section 3.1), a massive improvement is gained from using LM embeddings (Table 3, Figure 2).

**Figure 2:**
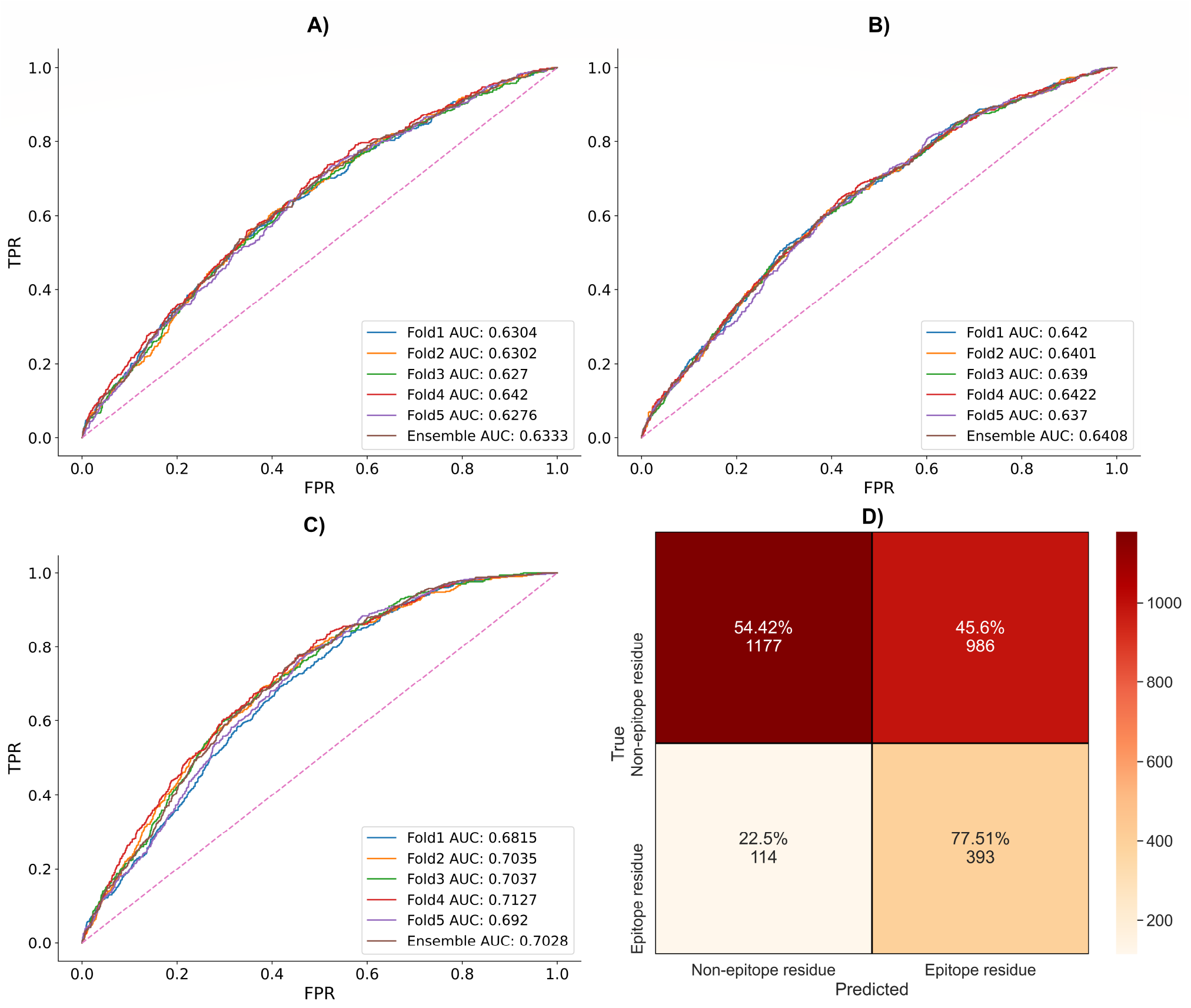
ROC-AUC curves for the BP3C50ID FFNN, illustrate the difference of using sparse (**A**), BLOSUM62 (**B**) or ESM-1b encodings (**C**). The x and y axis are the false and true positive rates respectively. Dashed lines along the diagonal indicate random performance at 50 % AUC, and the remaining lines are the performances of different fold models. Also, a confusion matrix illustrates threshold-dependent performance of the best FFNN (ESM-1b) model (**D**). The true negatives or positives and predicted negatives or positives are on the vertical and horizontal axis respectively.

### 3.3 Feature engineering: Adding sequence lengths improves performance

In BepiPred-2.0, one of the main factors that contributed to predictive performance was relative solvent accessibility (RSA), an input feature predicted from the antigen sequence using NetSurfP-2.0 [13]. We uncovered a positive correlation between NetSurfP-2.0 predicted RSA values and BepiPred-3.0 B-cell epitope probability scores, indicating that information on solvent accessibility is, at least in part, encoded in the ESM-1b embeddings (see supplementary results section 6.3). To quantify to what extent RSA contributes to epitope predictions in BepiPred-3.0, we compared the performance of an FFNN trained on the BP3C50ID dataset with and without NetSurfP-3.0 RSA added as an input feature [8]. We evaluated the performance on the same external test set of 15 antigens. Here, we found that the added RSA feature failed to improve performance further, indicating that BepiPred-3 already considers this information in the ESM embeddings (Table 4). We note that NetSurfP-3.0 itself uses ESM-1b to predict residue RSA.

**Table 4:** A FFNN was cross-validated on BP3C50ID ESM-1b encodings as well as corresponding sequence lengths or NetSurfP-3.0 computed RSA values. The best models in terms of validation CE loss and AUC were evaluated on the external BP3C50ID test set of 15 antigens.

| Models | AUC | AUC10 | MCC |
| --- | --- | --- | --- |
| FFNN (+ NetSurfP-3.0 RSA) | 0.692 | 0.131 | 0.223 |
| FFNN (+ SeqLen) | 0.714 | 0.143 | 0.241 |

We also uncovered a negative relationship between the length of the antigen sequences and their respective epitope residue ratios, with a Spearman correlation coefficient of -0.58 (see supplementary results section 6.4). We expect this to be due to multiple effects, concerning the larger surface-to-volume ratio of small proteins, the issue of a limited number of antibodies being mapped to individual antigens, as well as possible selection biases in the dataset. Interestingly, we also found that the performance of our best models decreased for longer antigen sequences, suggesting that the models could not infer the protein sequence length from the sequence embeddings. To account for this trend, we trained another FFNN on the BP3C50ID dataset, where sequence lengths were added as an additional input, and the resulting models evaluated on the same 15 antigen test set. This further improved our AUC performance from 0.697 to 0.714, and in a paired t-test comparing both models CE loss scores on all test antigen residues, we determined that the performance increase was statistically significant with a p-value of < 1 × 10^*−*10^. We therefore used this as the final model for the BepiPred-3.0 web server and for further benchmarking (sections 3.4 and 3.5).

### 3.4 BepiPred-3.0 web server

BepiPred-3.0 is an easy to use tool for B-cell epitope prediction, as the user only needs to upload protein sequence(s) in fasta format. Furthermore, one can specify the threshold for epitope classification, and by default, a threshold of 0.1512 is used. Alternatively, the user may specify the number of top residue candidates that should be included for each protein sequence. These options generate two separate fasta formatted output files.

A total of 4 result files are generated. One is a fasta file containing epitope predictions at the set threshold, and another is a similar fasta file, but epitope predictions are instead the number of top residue candidates specified by the user. In each file, epitope predictions are indicated with letter capitalization. The third is a .csv file containing all model probability outputs on the uploaded protein sequence(s). Finally, we provide a .html file, which works as a graphical user interface that can be opened in any browser. Similar to BepiPred2.0, this interface can be used for setting a threshold for each antigen and downloading the corresponding B-cell epitope predictions [9]. Due to memory limitations, however, this interface is limited to the first 30 sequences in the uploaded fasta file. We believe this intuitive interface will allow researchers to maximize their precision of B-cell epitope prediction, as a single threshold might not work for all uploaded sequences (Figure 3).

**Figure 3:**
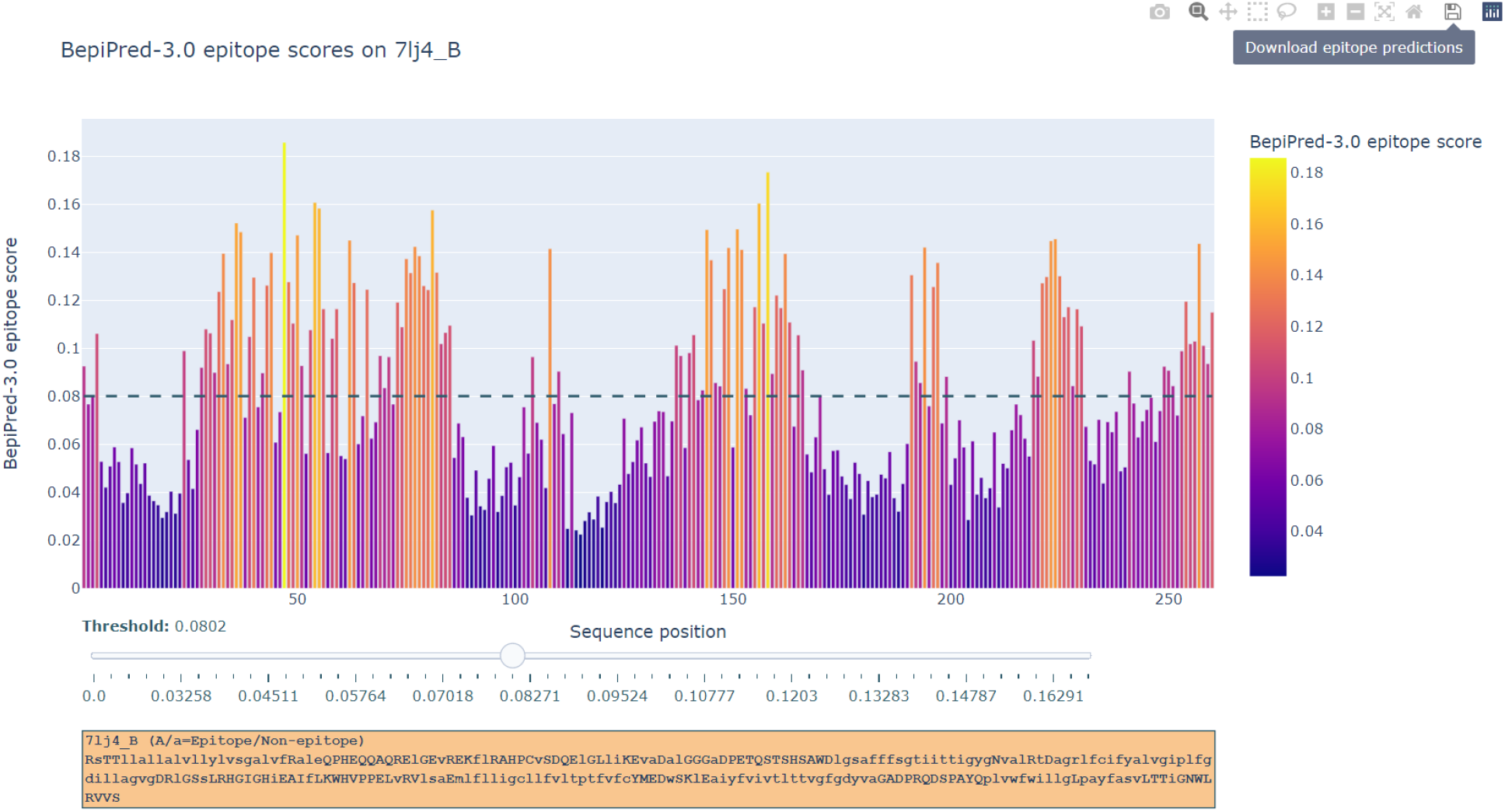
The graphical user interface for BepiPred-3.0 on the external test set protein 7lj4_B. In this interface, the x and y axis are protein sequence positions and BepiPred-3.0 epitope scores. Residues with a higher score are more likely to be part of a B-cell epitope. The threshold can be set by using the slider bar, which moves a dashed line along the y-axis. Epitope predictions are updated accordingly, and B-cell epitope predictions at the set threshold can be downloaded by clicking the button ‘Download epitope prediction’.

### 3.5 Benchmarking: BepiPred-3.0 outperforms its predecessors as well as structure-dependent B-cell epitope prediction tools

BepiPred-3.0 was re-evaluated and compared to its two predecessors on the 5 antigens from the BepiPred-2.0 paper external test set for a direct benchmarking [9][7]. Here, we find a drastic improvement in BepiPred 3’s AUC performance versus its predecessors, at 0.57, 0.60 and 0.71 for BepiPred 1, 2 and 3, respectively (Table 5).

**Table 5:** Benchmarking on 5 antigen external test set from BepiPred2 paper.

| Models | AUC | AUC10 | Data | Method |
| --- | --- | --- | --- | --- |
| BepiPred-1.0 | 0.573 | 0.055 | Peptides | HMM |
| BepiPred-2.0 | 0.596 | 0.080 | PDB | RFC |
| BepiPred-3.0 | **0.710** | **0.129** | PDB | NLP |

When tested on the IEDB external test set (see method section 2.3), BepiPred-3.0 obtained an AUC score of 0.65 when using ESM-1b sequence embeddings, versus 0.50 if using either sparse or BLOSUM62 encodings [27]. This demonstrates the improved ability to generalize on novel datasets when using the ESM-1b protein language model embeddings (Table 6).

**Table 6:** Benchmarking on homology reduced IEDB dataset of 3560 sequences.

| Models | AUC | AUC10 |
| --- | --- | --- |
| BepiPred-3.0 (Sparse) | 0.497 | 0.041 |
| BepiPred-3.0 (Blosum62) | 0.501 | 0.043 |
| BepiPred-3.0 (ESM-1b) | **0.644** | **0.148** |

We also benchmarked against a recently developed structure-based B-cell epitope predictive tool, epitope3d, that was in turn shown to outperform different other tools [6][29][16][26]. A 5-fold cross-validation setup on the 200 antigens available at the epitope3D online tool, was used for re-training and validating Bepipred-3.0. We then evaluated the re-trained model on the provided epitope3d external test set composed of 45 antigens. The evaluations in the epitope3D paper were done only on surface residues, and so to ensure a fair comparison, we calculated BepiPred’s performance on a subset of surface residues with an RSA above 15%, as defined in the epitope3D paper..

While epitope3D obtains an AUC Of 0.59 on their test set, BepiPred-3.0 obtains an AUC of 0.7, when evaluated on surface residues only (Table 7). We also tried including non-surface residues in the evaluation as well, which improved the AUC score to 0.74. We find these results surprising, as structure-based methods are generally considered superior to sequence-based methods, indicating the sheer power of the recent developments in protein language modeling.

**Table 7:** Benchmarking on surface residues of the epitope3D external test set of 45 antigens. Performance numbers for epitope3D were extracted from the publication.

| Models | AUC surface only |
| --- | --- |
| BepiPred-3.0 | **0.70** |
| Epitope-3d | 0.59 |
| Seppa-3.0 | 0.52 |
| Discotope | 0.49 |
| Ellipro | 0.44 |

## 4 Discussion

Deep learning methods, such as ESM-1b and AlphaFold, are revolutionizing the field of biology at large, and changing the role of computational tools in numerous tasks [17][10]. In this paper, we demonstrate that protein LMs can vastly improve B cell epitope prediction, and, using only the antigen sequence as an input, outperform existing tools, including structure-based ones. We can envision that, by using a similar approach on structure based embeddings calculated on solved antigen structures, or on structural models created using AlphaFold, it will be possible to further improve the current results [23][10]. We also want to argue on the fact that the current BepiPred-3.0 results are likely affected by the limited availability of experimental structures. The available solved antibody-antigen complexes are just a minute fraction of all possible pathogenic proteins, and of the antibodies that target them. Due to the underrepresentation of observed epitopes in current datasets, we expect that in many cases, regions predicted to be epitopes may not be false positives, but rather should be considered unlabeled or potentially positive residues due to data paucity. To this aim, it is possible to frame the epitope prediction problem as a positive and unlabeled (PU) training problem. Moreover, we can also argue that the current AUC of the model, around 71%, is an underestimation, and only by collecting more experimental data will it be possible to fully assess how close we are to the upper limit of the B cell epitope prediction tools.

It is also interesting to note that a major progress in this class of predictors would be the possibility to include the sequences of individual antibodies, or of antibody libraries, for which we want to identify all the potential epitopes. Language models provide an elegant way to include them in the prediction, by encoding the antibodies together with the antigen. As more data will be available, it will be interesting to test if LMs can also provide a solution to this fundamental problem in immunology and biotechnology.

To conclude, BepiPred-3.0 is available as a web server and as a stand alone software, it is easy to use for experts and non-experts alike, and provides state-of-the art B cell epitope predictions that will be fundamental to tasks of primary medical and societal importance, such as vaccine development and antibody engineering.

## Supporting information

All supplementary methods and results

