## Supplementary material for "BepiPred-3.0: Improved B-cell epitope prediction using protein language models": All supplementary methods and results

### 5 Supplementary methods

#### 5.1 BP3NC dataset

The epitope collapse strategy can result in sequence insertions. This in particular for the BP3C50ID data set. Therefore, models trained on the BP3C50ID dataset may not generalize as well on non-collapsed antigens. To investigate this, a dataset consisting of ESM-1b encodings from non-collapsed sequences was constructed [17]. The 1466 BP3 sequences were clustered at 20% sequence identity, as calculated by MMseqs2, and the 15 test and other antigens of the same respective clusters were placed in an external test set [22]. The remaining sequences were clustered at a 70% identity, and put into 5 partitions. For each cluster, multiple sequence alignments (MSA) were made using a local installment of the MUSCLE multiple sequence alignment tool [5]. ESM-1b encodings were then picked from the alignments: If any residue was positively annotated, an ESM-1b encoding was randomly picked from a sequence, where it was annotated as part of an epitope. If there were no positives at a position, a random ESM-1b encoding was picked from a sequence, that was not a gap. Corresponding sequence lengths were also concatenated to each ESM-1b encoding. The resulting number of positional ESM-1b encodings for each of the partitions were 19345, 18340, 18373, 15907 and 19730 respectively. For the external test set, 2615 positional ESM-1b encodings were obtained. The dataset was named BepiPred3 non-collapsed (BP3NC).

#### 5.2 B-cell epitope prediction on BP2 and BP3 datasets with RFCs

RFCs were trained on 5-fold cross-validation setups of sparse or BLOSUM62 encodings on datasets BP2, BP2HR, BP3, BP3HR, BP3C50ID. During cross-validation, forest sizes ranging from 25-300 were tested and the optimal forest size chosen on a validation AUC score basis (Table 8). All models were evaluated on an external test dataset of five antigens, and the BP3 and BP3C50ID models were also evaluated on the external test set of 15 antigens, which served as a baseline for the deep learning models in this study.

**Table 8:** The optimal forest size for RFCs trained on BP2, BP2HRR, BP3, BP3HR and BP3C50ID. The tested forest size ranged from 25-300, and the optimal size chosen on a validation AUC score basis.

| Datasets | Fold1 | Fold2 | Fold3 | Fold4 | Fold5 |
|---|---|---|---|---|---|
| BP2 (Sparse) | 225 | 250 | 250 | 275 | 275 |
| BP2 (BLOSUM62 | 275 | 250 | 225 | 300 | 225 |
| BP2HR (Sparse) ) | 225 | 225 | 200 | 250 | 300 |
| BP2HR (BLOSUM62) | 175 | 225 | 225 | 125 | 125 |
| BP3 (Sparse) | 125 | 250 | 175 | 275 | 275 |
| BP3 (BLOSUM62) | 300 | 225 | 300 | 275 | 250 |
| BP3HR (Sparse) | 250 | 225 | 250 | 275 | 225 |
| BP3HR (BLOSUM62) | 225 | 200 | 300 | 300 | 250 |
| BP3C50ID (Sparse) | 225 | 225 | 175 | 225 | 300 |
| BP3C50ID (BLOSUM62) | 275 | 275 | 275 | 250 | 300 |

#### 5.3 B-cell epitope prediction on BP3 datasets with FFNN, CNN and LSTM

FFNNs, CNNs and LSTMs were trained on BP3 and BP3C5ID using sparse, BLOSUM62 or ESM-1b encodings. All models underwent hyperparameter tuning, and the best model was chosen by CE loss score averaged across all validation splits. The hyperparameter tuning varied over learning rates, weight decays, dropout rates and different architectural setups. The FFNN consisted of an input layer with 1280 or 189 nodes depending on whether ESM-1b or BLOSUM62 and sparse encodings were used. Each residue input was passed through 3 linear layers, where the number of nodes varied for different datasets and encodings. Between the input and first layer, first and second layer, second and third layer, ReLU activation and varying degrees of dropout were used. The output from these layers was passed to a final layer consisting of 2 nodes, whereupon a softmax function mapped each node output to a class probability. Finally, cross entropy loss was used to obtain classification loss scores, which were backpropagated through the network. All model weights, including the LSTM and CNN, were updated using an Adam optimizer with varying degrees of learning rate and weight decay (Table 9).

The CNN consisted of 2 1-dimensional convolutional layers with different kernel sizes. The input dimension to the convolutional layers was (batch size, 1280 or 189, longest sequence length), and each sequence was padded according to the longest sequence in the batch. A small batch size of 4 was used for the FFNN, CNN and LSTM. The kernel for each convolutional layer convolutes across sequence positions of each antigen in the batch. This can be done with different number of kernels, also sometimes referred to as the output channel. Each position is

**Table 9:** FFNNs were trained on datasets BP3 and BP3C50ID using sparse, BLOSUM62 or ESM-1b encodings. For each setting, different architectures and training parameters were tested. The best model was chosen on CE loss score averaged across validation splits.

| FFNN hyperparameters | LR | Weight decay | Layers | Dropout |
|---|---|---|---|---|
| BP3 (Sparse) | 0.0001 | 0.01 | [90, 45, 30] | [0.75,0.75,0.6] |
| BP3 (BLOSUM62) | 0.0005 | 0.01 | [150,120,45] | [0.75,0.75,0.6] |
| BP3 (ESM-1b) | 0.0001 | 0.01 | [180,90,45] | [0.7,0.7,0.7] |
| BP3C50ID (Sparse) | 0.0001 | 0.005 | [150,120,64] | [0.7,0.7,0.7] |
| BP3C50ID (BLOSUM62) | 0.0001 | 0.005 | [150,120,64] | [0.75,0.75,0.6] |
| BP3C50ID (ESM-1b) | 0.0001 | 0.005 | [180,90,45] | [0.7,0.7,0.7] |

mapped to an output that matches the specified number of kernels. These outputs were then passed through max pooling layers with stride 4, whereafter each positional output was passed through a final 3-layer densenet consisting of 64, 32 and 2 nodes. The classification error, backpropagation and weight updating was done in a similar fashion as the FFNN (Table 10).

**Table 10:** CNNs were trained on datasets BP3 and BP3C50ID using sparse, BLOSUM62 or ESM encodings. For each setting, different architectures and training parameters were tested. The best model was chosen on CE loss score averaged across validation splits. A 3-layered FFNN consisting 64, 32 and 2 nodes was put atop of all CNNs.

| CNN hyperparameters | LR | Weight decay | Nr. of kernels | Dropout | Kernel |
|---|---|---|---|---|---|
| BP3 (Sparse) | 0.0001 | 0.005 | 108 | [0.75,0.75] | [3,5] |
| BP3(BLOSUM62) | 0.0005 | 0.01 | 108 | [0.7,0.7] | [3,5] |
| BP3 (ESM-1b) | 0.001 | 0.005 | 108 | [0.7,0.7] | [3,5] |
| BP3C50ID (Sparse) | 0.0001 | 0.005 | 108 | [0.7,0.7] | [3,5] |
| BP3C50ID (BLOSUM62) | 0.0005 | 0.01 | 108 | [0.75, 0.75] | [5,7] |
| BP3C50ID (ESM-1b) | 0.0001 | 0.01 | 112 | [0.75,0.75] | [1,3] |

When working with ESM-1b encodings, a 3-layered FFNN consising of 1280, 680 and 128 nodes was added before a bidirectional LSTM. Between the input and first layer, ReLU activation and a dropout rate of 0.7 was used. This output was fed through a bidirectional single layered LSTM with input dimension (batch size, sequence length, 128) and a hidden state size of 50. As the LSTM was bidirectional, a total of 100 hidden states were obtained for each position in a sequence. Each positions hidden state was fed through a FFNN with an input layer of size 100, and 3 layers consisting of 64, 32 and 2 nodes. Between the input and first layer, first and second layer, ReLU activation and dropout rates of 0.75 were used.
For the BLOSUM62 and sparse encodings, a bidirectional single layered LSTM with input dimension (batch size, sequence length, 189) and hidden state of 100 was used. Each positions hidden state was then fed through a densenet with an input layer size of 200 and 3 layers consisting 120, 48 and 2 nodes. Between the input and first layer, first and second layer, ReLU activation and dropout rates of 0.7 were used. A learning rate of 0.0001, weight decay of 0.005 and an Adam optimizer was used when training both LSTMs. An extensive hyperparameter tuning was done for the LSTM, experimenting with different LSTM layer sizes, dropout rates between LSTM layers, varying dropout rates for the dense networks before and after the LSTM, the weight decay values and the hidden state sizes. In the end, using single-layered LSTMs and adding high dropout rate before and after LSTM had performance on par with multi-layered LSTMs.

After having found optimal hyperparameters for above-mentioned models, they were re-trained on their respective dataset. A model for each cross-validaiton fold was obtained by choosing the ones that yielded the best validation CE loss and AUC scores.

Antigen ESM-1b encodings were also combined with corresponding sequence lengths or NetSurfP-3.0 RSA values. These experiments were only done for FFNNs using the same architecture and training parameters as when training BP3C50ID FFNN ESM-1b models in Table 9, except the input layer was now set to 1281 or 1282 and all dropout rates to 0.65.

Finally, a FFNN with input layer of size 1281 was trained on ESM-1b encodings + sequence lengths of the BP3NC dataset. The same architecture and training parameters as the BP3 FFNN ESM-1b model were used (Table 9).

### 6 Supplementary results

#### 6.1 Establishing a dataset: Random forest classifiers on BP3 datasets outperform those trained on BP2 datasets

RFCs were cross-validated on datasets BP2, BP2HR, BP3, BP3HR and BP3C50ID. For each dataset, fold models were evaluated in terms of AUC, AUC10, MCC, recall, precision, F1-score and accuracy on an external test set consisting of the 5 antigens from the BepiPred-2.0 paper [9].

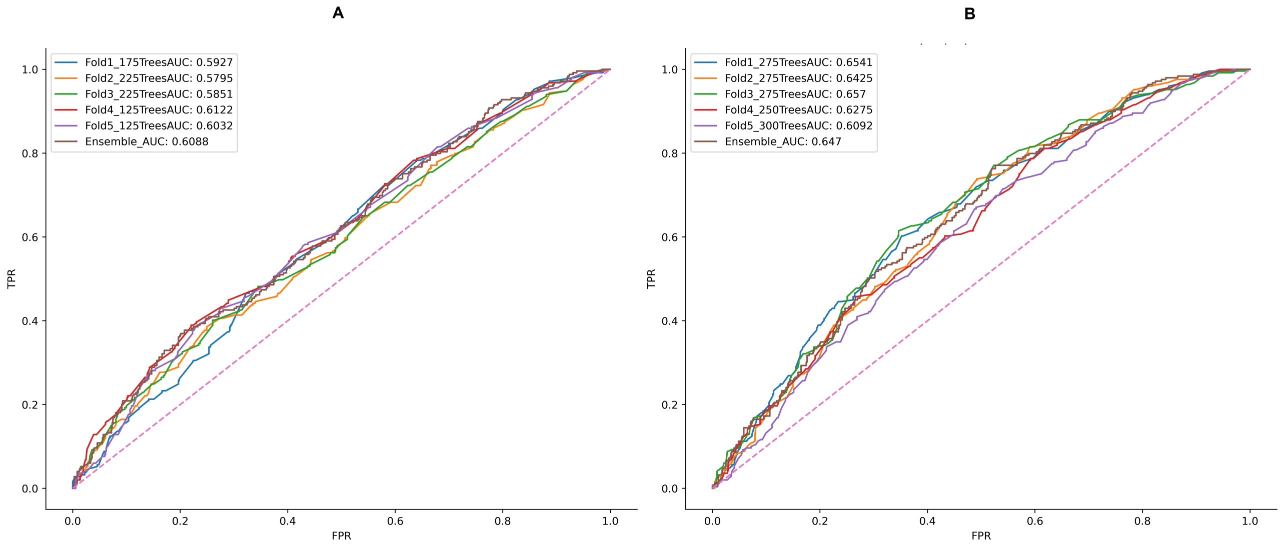


**Figure 4:** A ROC curve comparison of BP2HR (BLOSUM62) (**A**) and BP3C50ID (BLOSUM62) (**B**) on the 5 antigen external test set. The x and y axis are the false and true positive rates respectively (FPR and TPR). Dashed lines along the diagonal indicates a completely random performance, and the remaining lines are fold model performances.

**Table 11:** RFCs were cross-validated on sparse or BLOSUM62 encodings of datasets BP2, BP2HR, BP3, BP3HR and BP3C50ID. The best models in terms of validation AUC were evaluated on the 5 antigen external test set. The evaluative metrics used were AUC, AUC10, MCC, recall, precision, F1-score and accuracy. The best score within a metric category is marked in bold.

| Models | AUC | AUC10 | MCC | Recall | Precision | F1 | Accuracy |
|---|---|---|---|---|---|---|---|
| BP2 (Sparse) | 0.565 | 0.072 | 0.015 | 0.002 | **0.429** | 0.005 | **0.743** |
| BP2 (BLOSUM62) | 0.599 | 0.081 | 0.042 | 0.080 | 0.328 | 0.128 | 0.722 |
| BP2HR (Sparse) | 0.568 | 0.084 | 0.062 | 0.418 | 0.292 | 0.344 | 0.591 |
| BP2HR (BLOSUM62) | 0.594 | 0.098 | 0.097 | 0.274 | 0.34 | 0.303 | 0.677 |
| BP3 (Sparse) | 0.629 | 0.100 | 0.114 | 0.364 | 0.337 | 0.35 | 0.653 |
| BP3 (BLOSUM62) | 0.633 | 0.097 | 0.102 | 0.31 | 0.337 | 0.323 | 0.667 |
| BP3HR (Sparse) | 0.619 | 0.087 | 0.157 | 0.542 | 0.339 | 0.417 | 0.611 |
| BP3HR (BLOSUM62) | 0.63 | **0.105** | 0.133 | 0.471 | 0.334 | 0.391 | 0.623 |
| BP3C50ID (Sparse) | 0.616 | **0.105** | 0.143 | 0.52 | 0.332 | 0.406 | 0.609 |
| BP3C50ID (BLOSUM62) | **0.638** | 0.087 | **0.169** | **0.566** | 0.342 | **0.427** | 0.61 |

Models trained on BP3 datasets (BP3, BP3HR or BP3C50ID) compared to those trained on BP2 datasets (BP2 or BP2HR), have better performance in almost all evaluative metrics, and most importantly in terms of AUC and MCC. The best evaluation metric scores are marked in bold, and with the exception of accuracy and precision, all bold markings are scored by models from BP3 datasets. This performance boost is perhaps not surprising, as BP3 datasets incorporate more antigens. We determined that the BP3 (BLOSUM62) model performance increase over the BP2 (BLOSUM62) model was statistically significant at all common thresholds with a p-value of 1 x $10^{-15}$. The paired t-test was performed by comparing CE loss scores of both models evaluated on all test antigen residues. The use of BLOSUM62 encodings instead of sparse encodings increases AUC, and it also seems to increase MCC scores with the exception of BP3 and BP3HR. BLOSUM62 encodings contain information about amino acid similarity, and are more informative than sparse encodings, making this a nice display of how biological information can contribute to model performance. The epitope collapse strategy used for the BP3C50ID dataset enhances B-cell epitope predictions, as BP3C50ID (BLOSUM62) obtains the best AUC and MCC performance of 0.638 and 0.169 (Table 11). In a similar paired t-test as the one done above, we determined that the BP3C50ID (BLOSUM62) performance increase over BP3 (BLOSUM62) was statistically significant at a p-value of 1 x $10^{-20}$. Homology reduction impacts the MCC, but there isn't a consistent AUC performance increase overall. However, as the BP3C50ID dataset is the most homology reduced dataset, and the BP3C50ID (BLOSUM62) models boast the best scores in terms of AUC, MCC, positive recall or F1-score, then it is not far-fetched to say that it likely has an effect (Table 11).

A ROC curve comparison, shows that BP3C50ID (BLOSUM62) is clearly superior to BP2HR (BLOSUM62) as it performs best for all folds and the ensemble model (Figure 4).

#### 6.2 Improved performance using neural networks and ESM-1b sequence embeddings: BP3 dataset

**Table 12:** Evaluation performance on an external test set of 15 antigens, with a set of RFC, FFNN, CNN and LSTM models, trained using cross-validation on the BP3 dataset. The best models in terms of validation CE loss and AUC score were chosen for evaluation on the test set (see methods section 2.6). The evaluative metrics used were AUC, AUC10, MCC, recall, precision, F1-score and accuracy. The best score within a metric category is marked in bold. We also computed paired t-test p-values, comparing the CE loss scores of models using BLOSUM62 encoding and ESM-1b embeddings as input.

| BP3 Models | AUC | AUC10 | MCC | Recall | Precision | F1 | Accuracy | P-value |
|---|---|---|---|---|---|---|---|---|
| RFC (Sparse) | 0.604 | 0.096 | 0.074 | 0.356 | 0.236 | 0.284 | 0.658 | - |
| RFC (BLOSUM62) | 0.604 | 0.088 | 0.072 | 0.324 | 0.237 | 0.274 | **0.674** | - |
| FFNN (Sparse) | 0.617 | 0.108 | 0.132 | 0.564 | 0.250 | 0.346 | 0.595 | - |
| FFNN (BLOSUM62) | 0.637 | 0.109 | 0.163 | 0.620 | 0.260 | 0.366 | 0.593 | - |
| FFNN (ESM-1b) | **0.677** | **0.116** | **0.184** | 0.610 | **0.275** | **0.379** | 0.620 | 1 x $10^{-16}$ |
| CNN (Sparse) | 0.629 | 0.104 | 0.145 | 0.577 | 0.255 | 0.354 | 0.600 | - |
| CNN (BLOSUM62) | 0.629 | 0.070 | 0.157 | 0.618 | 0.257 | 0.363 | 0.588 | - |
| CNN (ESM-1b) | 0.664 | 0.102 | 0.180 | 0.535 | 0.284 | 0.371 | 0.656 | 1 x $10^{-5}$ |
| LSTM (Sparse) | 0.609 | 0.087 | 0.113 | 0.634 | 0.233 | 0.340 | 0.534 | - |
| LSTM (BLOSUM62) | 0.635 | 0.101 | 0.149 | **0.677** | 0.245 | 0.360 | 0.543 | - |
| LSTM (ESM-1b) | 0.652 | 0.103 | 0.165 | 0.551 | 0.272 | 0.364 | 0.634 | 1 x $10^{-9}$ |
| BP3 overall average | 0.632 | 0.099 | 0.139 | 0.551 | 0.255 | 0.346 | 0.609 | |

To assess the best performing machine-learning architecture and sequence representation for the BP3 dataset, optimal hyperparameters for four different architectures (RFCs, FFNNs, CNNs and LSTMs) were identified using a grid search (see methods section 2.5). For each architecture, we investigated the performance when representing protein residues with sparse encoding, BLOSUM62 encoding or ESM-1b embedding. The best performing models were selected from optimal cross-validation performance, and tested on the 15 antigen external test set using a classification threshold optimized on the validation splits (Table 12). We found that all neural networks using ESM-1b sequence embeddings as input performed better. We determined that the performance increase of models using ESM-1b embeddings instead BLOSUM62 encodings was statistically significant at all common thresholds with p-values of 1 x $10^{-16}$, 1 x $10^{-5}$ and 1 x $10^{-9}$ for the FFNN, CNN and LSTM respectively. The paired t-test was performed by comparing each models CE loss scores on all test antigen residues.

#### 6.3 Epitope probability scores are correlated with NetSurfP-2.0 computed relative surface accessibilities (RSA)

RSA is important for B-cell epitope prediction, because if a residue is not on the antigens surface, then antibody interaction is less likely. It is therefore not far-fetched to hypothesize that residues with low RSA are generally more unlikely to be epitope residues, and vice versa. Furthermore in BepiPred-2.0, one of the factors that contributed to predictive performance was NetSurfP-2.0 computed RSA values. The models constructed in this paper instead rely on ESM-1b embeddings, and it would therefore be interesting to see if they contain some of the same information. This was investigated by plotting the RSA values of all residues from the 15 antigen external test set against corresponding epitope probabilities scores outputted by the BP3C50ID (ESM-1b+SeqLen) ensemble model. RSA values were computed by uploading the 15 antigen sequences to the NetSurfP-2.0 online tool, which among other things, computes RSA per residue [13].

In the 15 antigen external test set, there are 2163 non-epitope and 507 epitope residues respectively. From these data, the correlation between residue RSA values and positive epitope probability scores in terms of spearman and Pearson correlation coefficients was found to be 0.562 and 0.548 respectively (figure 5). An interesting observation

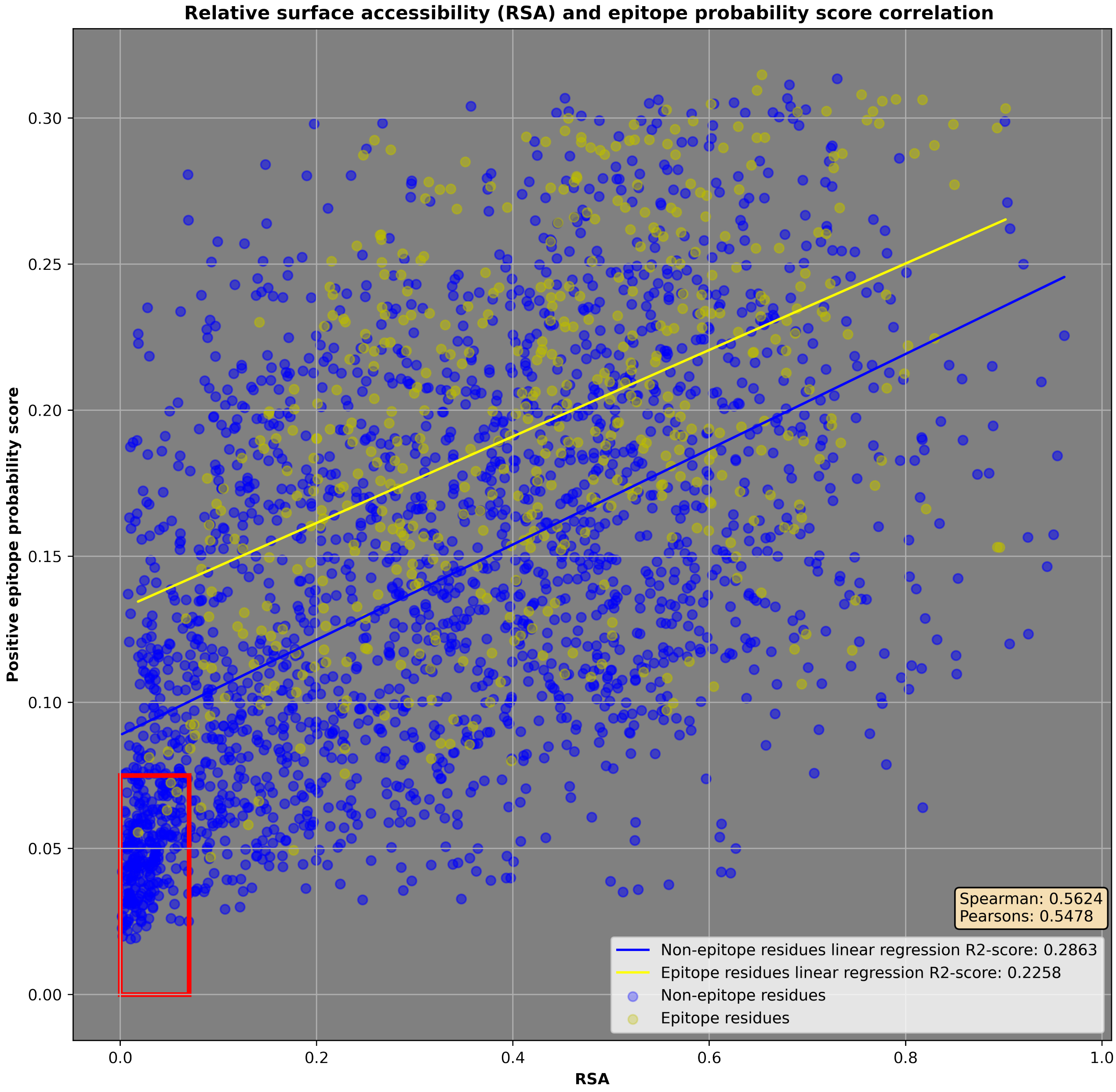


**Figure 5:** RSA values for amino acids (x-axis) plotted against corresponding epitope probability scores (y-axis) reveal a moderate positive correlation. These RSA values were computed using the NetSurfP-2.0 online tool on the 15 antigen external test set. Corresponding positive epitope probability scores were computed by the BP3C50ID (ESM-1b+SeqLen) ensemble model. Each dot corresponds to a residue, and blue or yellow indicate if it is part of an epitope or not. A red rectangle encapsulates an area dense with non epitope residues, stretching from 0.0-0.07 and 0.0-0.075 on the x- and y-axis. There are also two linear correlation plots for non-epitope (yellow) and epitope residues (blue). Spearman and pearson correlation coefficients comparing RSA and epitope probability scores was computed on all residues.

is the small area indicated by a red rectangle, which has a high density of non epitope residues receiving both low RSA and epitope probability scores. This area includes 265 non-epitope residues, which is approximately 0.123 of the total non-epitope residues. These residues suggest a group of easy to predict non-epitopes due to their very low RSA value. There are also 4 epitope residues in this group, but this accounts for less than 1% of the total epitope residues. In general, however, the plot indicates that epitope and non-epitope residues are difficult to distinguish. To indicate that they are distinguishable, two linear regressions for non-epitopes (blue) and epitope (yellow) residues are drawn. The epitope linear regression fitted line generally spans across an area of higher positive epitope probability scores. Also as one would expect, the average RSA value for epitope residues is higher than for non-epitope residues, 0.423 and 0.319 respectively. A paired t-test was also made for both RSA and epitope probability scores, to show that epitope and non-epitope residues constitute two different groups. The null hypothesis was that they were not two different groups. In both cases, it is statistically significant at all common thresholds with p-values of 1 x $10^{-40}$ and 1 x $10^{-109}$ when comparing RSA values and epitope probability scores respectively, suggesting that it is very unlikely that they are of the same group (Figure 5).

#### 6.4 Adding sequence lengths to ESM-1b encodings further improves performance

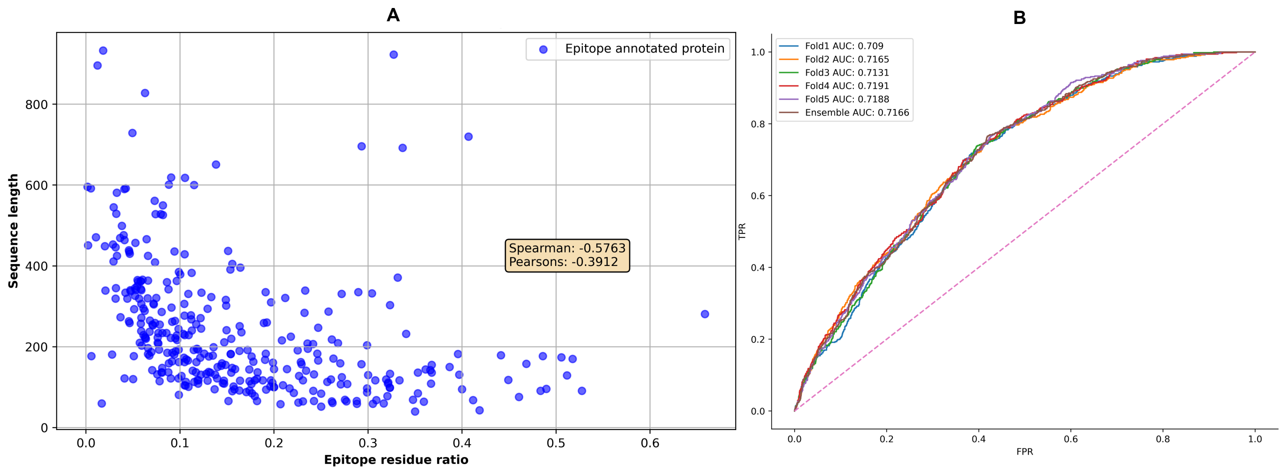


**Figure 6:** In the left figure (**A**), antigen epitope ratios (x-axis) plotted against corresponding sequence lengths (y-axis) uncover a moderate negative monotonic relationship with a spearman coefficient of -0.576 (left). In the right figure, a ROC curve of BP3C50ID FFNN (ESM-1b+SeqLen) performance on the 15 antigen external test set shows a performance boost compared to earlier constructed models (**B**).

Epitope residue ratios for antigens in the BP3C50ID dataset were computed as the total number of epitope residues divided by the total number of residues. These ratios were then plotted against corresponding sequence lengths, revealing a monotonic negative trend, indicating that smaller antigens tend to have relatively more epitope residues. Oppositely, this means that larger antigens have generally less epitope residues relative to their size. This trend is also indicated by a spearman correlation coefficient of -0.576, suggesting a moderate to strong monotonic relationship. This sprung the idea, that including sequence lengths as input could improve performance, as then the model could learn the relation between epitope residue ratio, and an antigens size (Figure 6).

#### 6.5 BP3C50ID models generalize on non-collapsed antigens

**Table 13:** Evaluation metrics on the BP3NC external test set show that models trained on collapsed antigens, generalize on non-collapsed antigens. The best score within a metric category is marked in bold.

| Models | AUC | AUC10 | MCC | Recall | Precision | F1 | Accuracy |
|---|---|---|---|---|---|---|---|
| BP3NC (ESM-1b+SeqLen) | **0.711** | 0.116 | 0.220 | 0.636 | **0.294** | 0.402 | **0.639** |
| BP3C50ID (ESM-1b+SeqLen) | 0.709 | **0.147** | **0.226** | **0.707** | 0.284 | **0.405** | 0.605 |

The BP3C50ID dataset has insertions resulting from collapsing antigens annotations at 50 % sequence identity, and a model trained on this dataset may therefore not generalize well on non-collapsed antigens. To investigate this, a new dataset called BP3NC was constructed using ESM-1b encodings from non-collapsed antigens (see supplementary method section 5.1). A FFNN was cross-validated on the BP3NC dataset and models with the best validation CE loss and AUC chosen. These models were evaluated on the BP3NC external test set, which was constructed from the 15 antigens external test set. For comparison and to investigate how well BP3C50ID models generalize on non-collapsed antigens, BP3C50ID (ESM-1b+SeqLen) was also evaluated on the test set.

The BP3NC (ESM-1b+SeqLen) models achieved AUC, AUC10 and MCC scores of 0.711, 0.116 and 0.220 respectively. The BP3C50ID FFNN (ESM-1b) had a similar performance of 0.709, 0.147 and 0.226 for AUC, AUC10 and MCC respectively. This is only slightly worse than what was observed earlier on collapsed antigens in Table 4, suggesting that it generalizes on non-collapsed antigens. It was expected that training on ESM-1b encodings from non-collapsed antigens could give the BP3NC FFNN model a performance boost above

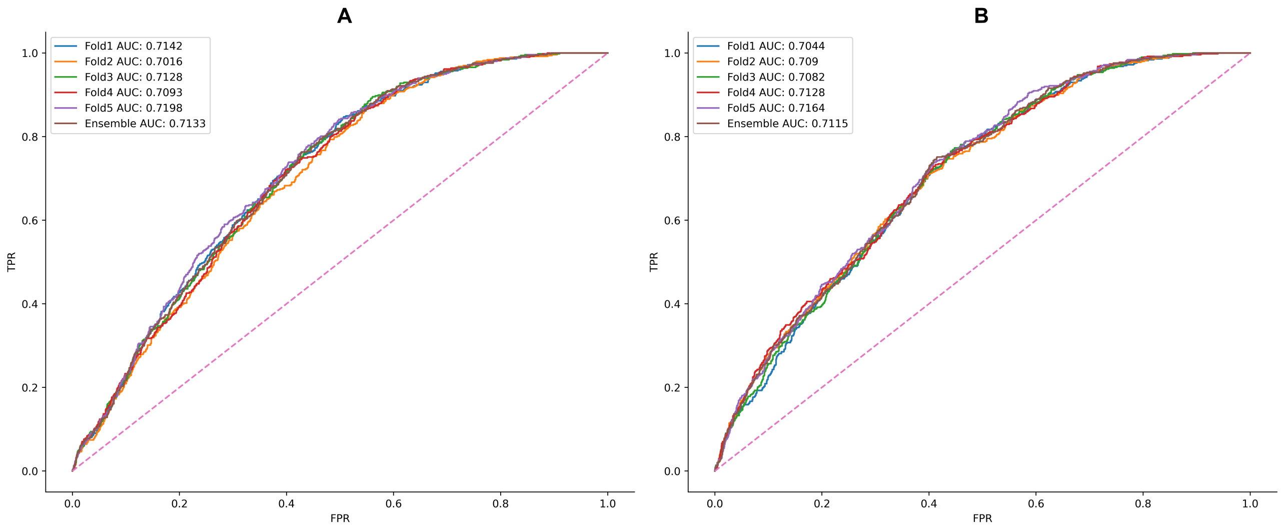


**Figure 7:** ROC curve comparisons of BP3NC (ESM-1b+SeqLen) (**A**) and BP3C50ID (ESM-1b+SeqLen) (**B**) on the BP3NC test set are similar, but differ slightly in the beginning and middle-end. The x and y axis are the false and true positive rates respectively (FPR and TPR). Dashed lines along the diagonal indicate a completely random performance, and the remaining lines are the performances of different fold models.

BP3C50ID (ESM-1b+SeqLen). However, they instead achieve very similar performance, indicating that collapsing annotations did not significantly affect how well the ESM1-b could realistically encode antigens (Table 13).
